# Exploring the Mosaic-like Tissue Architecture of Kidney Diseases Using Relation Equivariant Graph Neural Networks on Spatially Resolved Transcriptomics

**DOI:** 10.1101/2023.11.09.566479

**Authors:** Mauminah Raina, Hao Cheng, Hari Naga Sai Kiran Suryadevara, Treyden Stransfield, Dong Xu, Qin Ma, Michael T. Eadon, Juexin Wang

## Abstract

Emerging spatially resolved transcriptomics (SRT) technologies provide unprecedented opportunities to discover the spatial patterns of gene expression at the cellular or tissue levels. Currently, most existing computational tools on SRT are designed and tested on the ribbon-like brain cortex. Their present expressive power often makes it challenging to identify highly heterogeneous mosaic-like tissue architectures, such as tissues from kidney diseases. This demands heightened precision in discerning the cellular and morphological changes within renal tubules and their interstitial niches. We present an empowered graph deep learning framework, REGNN (Relation Equivariant Graph Neural Networks), for SRT data analyses on heterogeneous tissue structures. To increase expressive power in the SRT lattice using graph modeling, the proposed REGNN integrates equivariance to handle the rotational and translational symmetries of the spatial space, and Positional Encoding (PE) to identify and strengthen the relative spatial relations of the nodes uniformly distributed in the lattice. Our study finds that REGNN outperforms existing computational tools in identifying inherent mosaic-like heterogenous tissue architectures in kidney samples sourced from different kidney diseases using the 10X Visium platform. In case studies on acute kidney injury and chronic kidney diseases, the results identified by REGNN are also validated by experienced nephrology physicians. This proposed framework explores the expression patterns of highly heterogeneous tissues with an enhanced graph deep learning model, and paves the way to pinpoint underlying pathological mechanisms that contribute to the progression of complex diseases. REGNN is publicly available at https://github.com/Mraina99/REGNN.

## 1 Introduction

The kidneys play several vital roles in maintaining bodily equilibrium, including filtrating bodily fluids and waste, regulating blood acid-base balance, maintaining electrolyte balance, and supporting the production of red blood cells[1]. Among diseases that affect the kidney, Chronic Kidney Disease (CKD) and Acute Kidney Injury (AKI) are two of the most common diseases worldwide. In recent years, there has been a significant increase in the incidence of this disease, resulting in a global estimated CKD prevalence of approximately 13.4%. This has led to 5 to 7 million patients experiencing kidney failure in late-stage CKD[2]. Similarly, AKI was found to have a prevalence of up to 3,000 cases in 1 million hospitalized patients, and this rate tends to rise notably among those who are critically ill[3]. In the presence of AKI or CKD, acute and chronic cellular and morphological changes occur in renal tubules that surround the interstitial niche. Currently, there are still many unknowns in basic research and clinical practice in understanding the biological and pathological mechanisms of CKD and AKI, especially how different cells play different roles in key injury-related processes such as fibrosis, immune infiltration, and epithelial repair[4] in different kidney tissues.

The emergence of spatially resolved transcriptomics (SRT) has brought novel advancements and opportunities to uncover the fundamental pathogenesis behind a wide range of human diseases[5]. Adding the positional context of cell spots with corresponding high-throughput gene expression [6], the availability of SRT data enables novel analysis to reshape our understanding of cell spatial organization and their functional generation[7]. Equipped with SRT, the contemporary study of key processes and pathways in kidney molecular biology can be conducted with increased efficiency and precision, including cell-cell communications[8], spatially variable genes relating to spatial development[9, 10], and analysis of tissue architecture[11]. These insights on spatially distributed gene expression would be vital in interpreting the underlying biological and pathological processes in kidney tissues involved in CKD and AKI.

Currently, there is a plethora of computational methods under active development for identifying tissue architecture from SRT data. BayesSpace[12] uses statistical methods and Bayes inference to dissolve tissue architecture, and Giotto[13] utilizes graph-based clustering methods for spatial clustering. Emerging graph neural networks (GNNs)[14] achieve state-of-the-art performances in most graph-related tasks by learning low-dimensional representations through deep learning architectures, making them possible for modeling cell relations on SRT. SpaGCN[15] first uses deep learning to categorize the spatial domain based on graph neural networks, showing good performance when identifying cell spatial structures in brain cortex benchmark data[4]. RESEPT[16] adopts graph neural networks to learn low dimensional embeddings as RGB images and uses the ResNet50[17] model to process image segmentation as cell types. Besides gene expression information, SpaGCN[15] and SiGra[18] adopt the H&E images as additional information to improve the model performances. However, there are still huge gaps between existing computational methods and the need to robustly analyze, identify, and annotate diverse cell types in heterogeneous tissue structures, especially in kidney research.

In general, there are several key challenges to analyzing SRT data on kidney diseases. The first challenge is modeling the kidney’s heterogeneous, sparse, and mosaic-like cell types. Compared to other organs such as the brain cortex which has ribbon distributed regions, the major cell types in the kidney, i.e., epithelial, endothelial, and stromal cells, and their numerous functionally distinct cell sub-types are in close approximation to each other, organized in ∼1 million small nephron structures within each kidney[19]. **Figure 1** shows the distinctions between these two tissue types on 10X Visium SRT. In a pathologic section, kidney tubules are cut from different directions during the sample preparation, making the cell type distribution vary significantly within a region. This complexity poses a huge challenge for current computational modeling methods that are often trained and benchmarked on less heterogeneous ribbon-distributed tissue structures[4, 12]. The second challenge is the limited expressive power of graph neural networks. Classical GNNs use message passing, aggregating, and iteratively updating local neighbor information[20]. Message passing GNN is theoretically limited by the 1-order Weisfeiler-Lehman (1-WL) test[21] to distinguish if two given graphs are isomorphic or not[22]. Moreover, the nature of the regular topology in the lattice structures of the SRT makes it difficult to differentiate between nodes for they have similar topology in the modeled lattice graphs. The third challenge is how to efficiently annotate and interpret the vast omics datasets that are generated in a timely and cost-effective manner. Classical deep learning approaches usually rely on substantial amounts of well-labeled data to train their models. However, precisely linking underlying histology to SRT spots can take human pathologists weeks to annotate[23], which makes it expensive, time-consuming, and sometimes impossible to complete.

**Figure 1:**
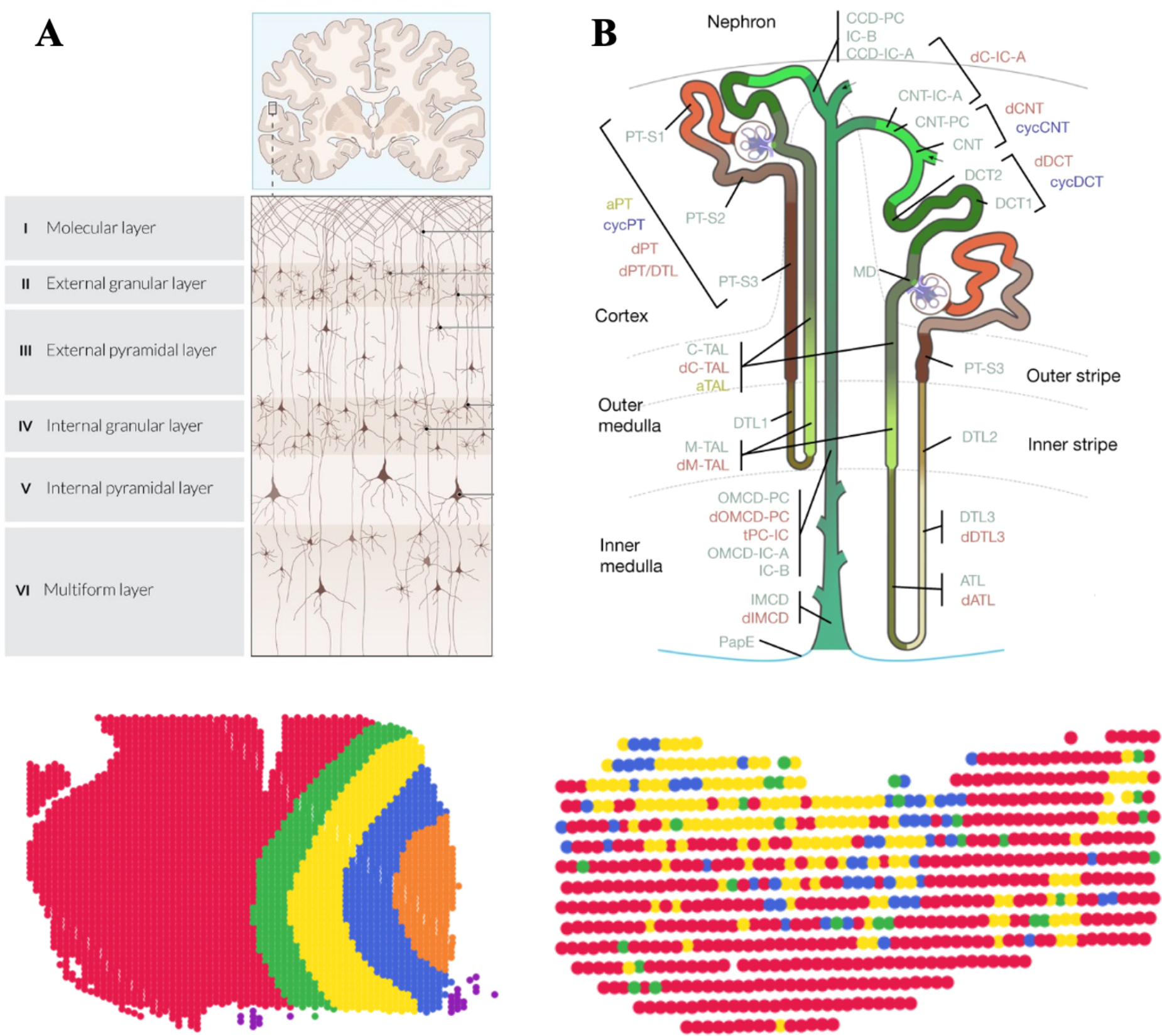
Comparison of the tissue architecture and cell type distribution of the Brain Cortex vs Kidney Nephron. A) Brain Cortex Diagram[25] and Ribbon-like Cell Types distribution of a brain cortex sample[26] on 10X Visium platform. B) Kidney Nephron Diagram[19] and Mosaic-like Cell type distribution of a kidney sample from KPMP[24] on 10X Visium platform. Each color represents a different cell type present in the tissue.

To address these challenges, we introduce an empowered graph deep learning framework, REGNN (Relation Equivariant Graph Neural Networks), for SRT data analyses on heterogeneous tissue structures. To increase the expressive power in the SRT lattice using graph modeling, the proposed REGNN integrates (1) equivariance to handle the rotational and translational symmetries of the spatial space, (2) Positional Encoding (PE) to identify and strengthen the relative spatial relations of the nodes uniformly distributed in the lattice. REGNN was tested on 23 kidney samples of 10X Visium SRT data from the KPMP (Kidney Precision Medicine Project) atlas [24], and REGNN outperformed existing competitive methods in various clustering-based criteria.

Our major contribution is designing an expressive power enhanced GNN framework to model heterogeneous SRT by improving aggregate and update operations in GNN’s message passing mechanism. Our contributions include: (1) The proposed REGNN integrates equivariance to model the symmetric nature of the SRT lattice in the aggregate operation. (2) REGNN integrates PE to mark the unique position of each graph node on the SRT lattice in the update operation. (3) REGNN is designed to address the mosaic-like heterogenous SRT data and achieves state-of-the-art performance on challenging kidney disease samples, which shows its potential in other highly heterogeneous tissues such as lymph nodes and colon.

## 2 Methods

We present REGNN, an empowered graph neural network designed for SRT data analyses on heterogeneous tissue structures. REGNN aims to capture the comprehensive relations between the spots in SRT by keeping their spatial relations equivariant and unique in the learned representation. Building on the classical GNN, REGNN incorporates two critical components to increase expressive power: equivariance and PE. The schema of the REGNN is shown in **Figure 2**. REGNN models the SRT as a graph, and it takes both expression and spatial coordinates as the input. This proposed model learns low-dimensional presentations of each spot of the SRT and infers tissue architectures from clustering the embeddings.

**Figure 2:**
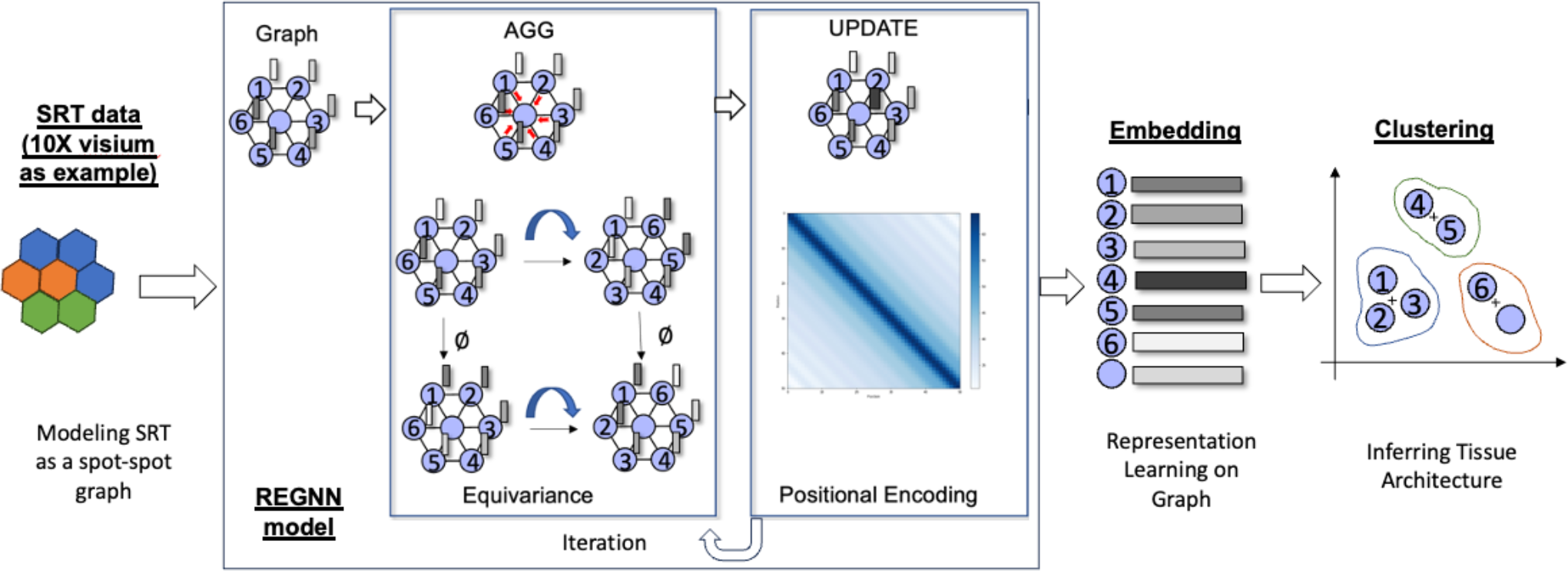
Schema of REGNN model. The empowered REGNN contains 1) Equivariance in AGG operation, and 2) Positional Encoding in UPDATE operation. Take 10X Visium platform as an example, REGNN models the SRT data as a spot-spot graph and learns the embeddings of the data, then inferred tissue architecture through clustering.

### 2.1 Graph modeling SRT and graph neural networks

Based on our in-house works RESEPT[16] and scGNN[14], SRT data are represented as a spatial spot-spot graph *G* = {*A, X*} by defining adjacency matrix *A, A* ∈ ℝ^|*V* |*×*|*V* |^ with node attributes *X, X* ∈ ℝ^|*V* |*×D*^, where *V* is the node set with *D* dimensions of gene expression. Each spot within a tissue sample containing a small number of cells is modeled as a node *ν*. The measured gene expression values of the spot are treated as the node attributes *X*, and the neighboring spots adjacent in the Euclidean space on the tissue slice are linked with an undirected edge *e*. As a result, this modeled undirected graph represents both the spatial context and the expression similarities between SRT nodes. This graph modeling method is applicable for both 10X Visium and MERFISH/smFISH platforms.

Generally, a classical *L*-layer graph convolution network (GCN) includes two steps of operations for each node *ν*_*i*_ at each layer: (1) AGG operation: aggregating messages 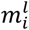 from the neighborhood at *l*-th layer as Eq (1); (2) UPDATE operation: updating node representation with 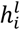 as Eq (2). The representation of each node 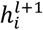 at layer *l* + 1 can be learned from the AGG and UPDATE operations.

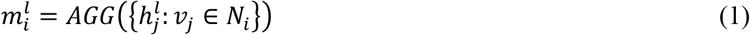

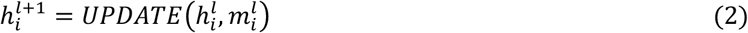

### 2.3 REGNN incorporates equivariance in AGG operation

For SRT in ST and 10X Visium technologies, directly modeling its lattice structure as a spot graph brings rotational and translational symmetries with the spatial gene expression pattern, which often confuses and diminishes the expressive power of classical GNNs. Inspired by E(n) equivariant GNN[27], REGNN integrates translation, rotation, reflection, and permutation equivariance with respect to an input set of spots in the modeled spatial spot-spot graph in the AGG operation. Targeting lattice symmetries, we define coordinates embedding *x* for each spot, where *x* ∈ ℝ^|*V*|*×*2^ in 2D SRT data. The initial *x*^0^ is the actual X-Y coordinates in the SRT lattice. Equivariance is integrated by modifying the classic GCN layer’s definition to include the learning of coordinates embeddings associated with each graph node[28]. For connected node *i* and *j* in the spot-spot graph at the *l*-th layer, REGNN defines message embedding *m*_*ij*_ in Eq (3) to incorporate the relative square distance between coordinates 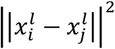, this information together with node embeddings 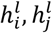, and edge attributes *e*_*ij*_ are summarized by learnable Multilayer Perceptrons (MLPs) *φ*_*e*_. Then coordinate embedding 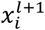 is updated by a weighted sum of all relative differences between coordinates with *m*_*ij*_ in Eq (4), where *φ*_*x*_ is another MLPs, *C* is a tunable hyperparameter to control the speed and strength. Compared with the classical AGG operation in Eq (1), REGNN updates the AGG operation by aggregating messages from all edges in Eq (5).

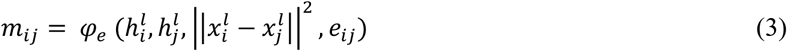

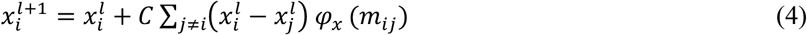

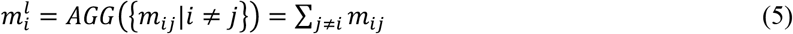

### 2.4 REGNN incorporates Positional Encoding in UPDATE operation

Besides integrating equivariance in handling symmetries in AGG operation, REGNN also incorporates PE in the following-up UPDATE operation. Widely used in linear data structures such as Transformers[29] and large language models[30], PE can often be treated as a unique feature to mark the entity’s relative position. In the SRT modeling, PE shows and strengthens the spatial relations between the nodes in the graph. Moreover, PE can be applied to the model to increase its discerning power to distinguish on an isomorphic graph. One widely accepted strategy in PE on the graph is to use sinusoids in the 2D space of the SRT[29, 31]. Thus, each coordinate is featured with a fixed PE consisting of *D* sinusoids with wavelengths that follow a geometric progression from 1 to the Nyquist limit, where *D* is the size of the SRT and *k* represents the coordinates of the SRT:

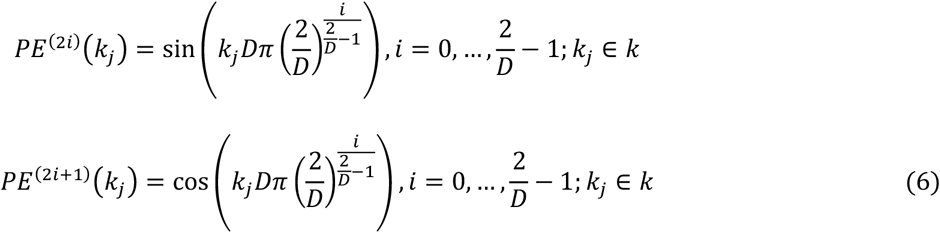

In REGNN, we update the UPDATE operation as Eq (2) in classical GNN by explicitly adding the PE embedding to message embedding 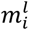 as Eq (7).

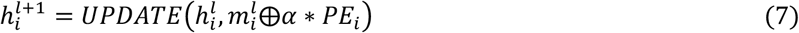

where *α* is the intensity of PE.

### 2.5 REGNN learns node embeddings in a graph autoencoder

Compared to classical *L*-layer GCN, REGNN uses the idea of equivariance in AGG operation as Eq (5) and integrates PE in UPDATE operation as Eq (7), the final formulation of REGNN is shown as Eq (8).

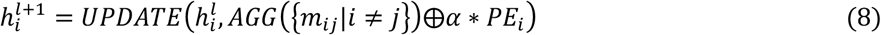

In SRT analysis, we use a graph autoencoder to learn node-wise low-dimensional representation on the modeled spatial spot-spot graph *A*. The proposed REGNN is utilized as the encoder of the graph autoencoder, and graph embedding *Z* is learned by stacking two layers of REGNN in Eq (9). The decoder calculates the inner product of *Z*, and then activated by sigmoid activation function to reconstruct the adjacency matrix *Â* in Eq (10).

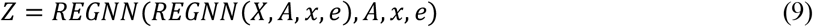

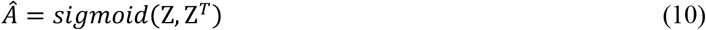

Overall, the loss function of the graph autoencoder is minimizing cross-entropy *L* between the reconstructed matrix *Â* and input adjacency matrix *A* as shown in Eq (11). With *N* nodes being the number of spots on the slide sample, *N* × *N* is the dimension of the adjacency matrix, and *a*_*ij*_ *and â*_*ij*_ are the elements of *A* and *Â* respectively.

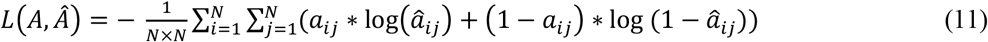

We assume the learned graph embedding *Z* represents the topological relations within the graph. Then k-means clustering algorithm is performed on *Z*, then the clustering results are annotated as distinct cell types in the tissue architecture.

REGNN was performed using different positional encoding intensity parameters, and various autoencoder dimensions for each test. The PE intensity parameters were 1.0, 1.5, 2.0, 2.5, 3.0, 3.5, 4.0, 4.5, and 5.0 while the autoencoder dimensions tested were 3, 8, 16, 32, 64, 128, and 256.

## 3 Results

### 3.1 REGNN identifies the tissue architecture of mosaic-like heterogenous kidney samples

In this study, 23 kidney samples sequenced by 10X Visium from healthy, CKD, and AKI patients were downloaded from the KPMP (Kidney Precision Medicine Project) atlas[24]. The sample information is detailed in **Supplementary Table 1**. The majority of cell types in the spot of the samples are annotated as epithelial cells, endothelial cells, immune cells, and stromal cells by experienced nephrology physicians from KPMP. These annotations are utilized as the gold standard benchmarks to test the performance of REGNN and other existing methods, including BayesSpace[12], Giotto[13], SpaGCN[15], RESEPT[16], and SiGra[18]. We used five criteria to quantify the efficacy of these SRT analysis tools, including Adjusted Rand Index (ARI), Rand Index (RI), Normalized Mutual Info score (NMI), Fowlkes Mallows Index (FMI), and Davies–Bouldin Index (DBI)[32].

Over all the healthy, CKD, and AKI samples, REGNN outperformed the competitive methods within a larger median and smaller deviation in ARI. The box plot of all these 23 samples is shown in **Figure 3**. Specifically, we can observe REGNN achieved better or comparable performance in ARI among 11 CKD samples (**Supplementary Figure 1**).

**Figure 3:**
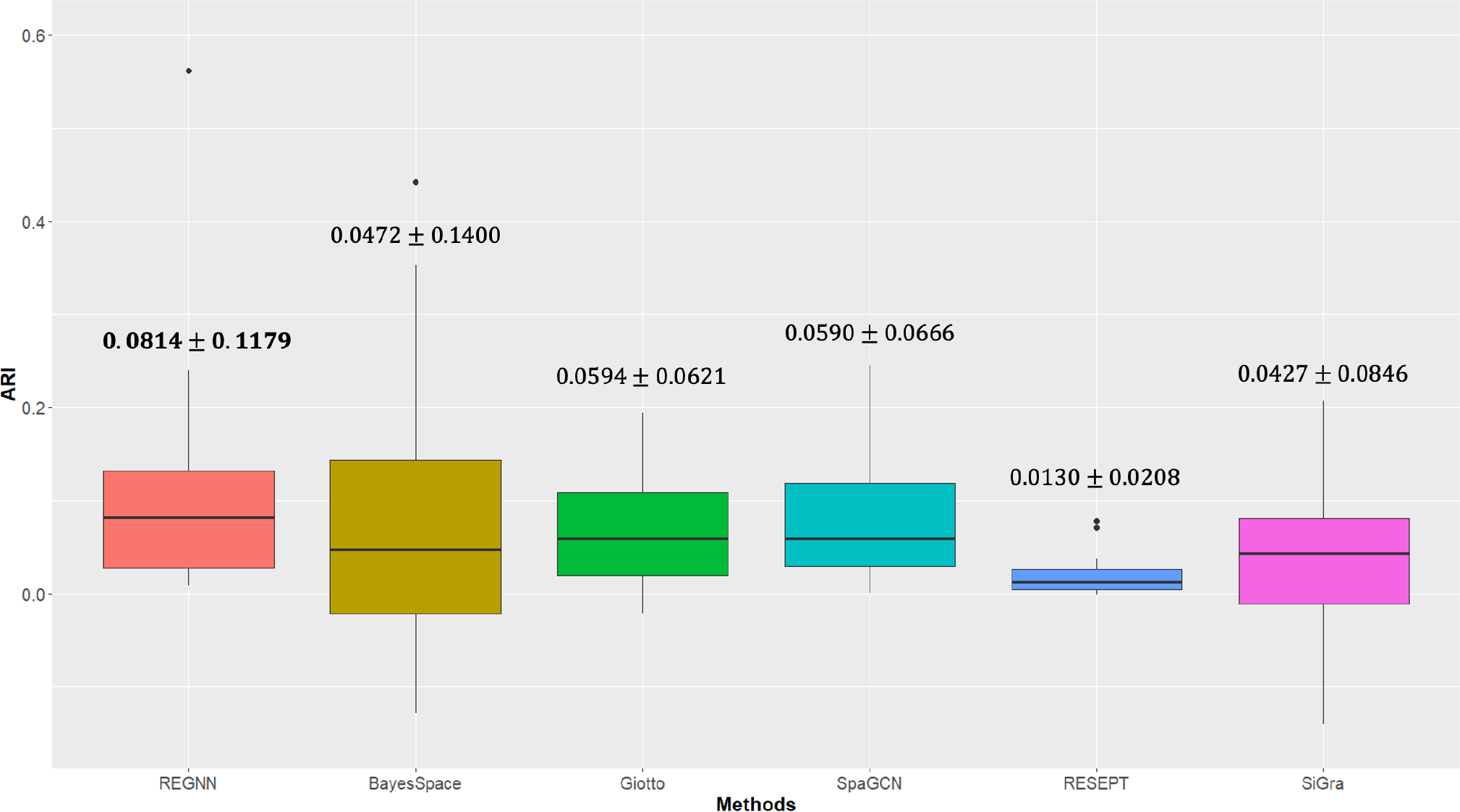
Performance comparison on ARI in all samples from KPMP. Bold for best results.

A success sample V10S14-085_XY04_21-0057 with a known presence of CKD was taken as a representative example in **Table 1**. In this sample, we observed REGNN leads the performance in nearly all five criteria with a clear margin. Especially, REGNN achieved 0.5814 in the ARI criteria, largely leading to the second-best BayesSpace’s 0.3535.

**Table 1:** Comprehensive performances on representative CKD V10S14-085_XY04_21-0057 sample.

| *ARI Performances* | ARI | NMI | RI | FMI | DBI |
| --- | --- | --- | --- | --- | --- |
| BayesSpace | 0.3535 | 0.4144 | 0.6618 | 0.6574 | **6.9389** |
| Giotto | 0.1938 | 0.3681 | 0.5625 | 0.5103 | 7.8053 |
| SpaGCN | 0.2456 | 0.3567 | 0.5938 | 0.5580 | 8.1301 |
| RESEPT | 0.0781 | 0.1155 | 0.5096 | 0.4663 | 9.8041 |
| SiGra | 0.2067 | 0.2822 | 0.5732 | 0.5321 | 10.850 |
| REGNN | **0.5814** | **0.4840** | **0.7875** | **0.8002** | 8.3409 |

Next, we scrutinized the computational methods by comparing their results to the gold standard annotations and mapping them to their original locations in **Figure 4**. On the representative CKD sample V10S14-085_XY04_21-0057, the epithelial cell and stroma cell populations are evenly spread, immune and endothelial cells are sparse and spread across the sample. Ideally, a computational method would be able to accurately identify these sparse cell groups while correctly categorizing adjacent spots as part of a larger cell-grouping. Some competitive methods such as RESEPT[16] and BayesSpace[12] only captured large homogeneous cell groups, while missing sparse cell groupings. Others like Giotto[13] and SpaGCN[15] were successful in identifying sparse spot clusters but were confused with their cell-type annotations. REGNN was able to correctly identify the larger sections of epithelial and stroma cells while also identifying some of the sparse spots that had been determined to be endothelial and immune cells. However, there were still some spots of immune cells that had been misclassified as epithelial, in addition to misclassifying a group of epithelial cells that had been localized within a section of stroma cells. Though REGNN shows a notable improvement over existing methods in identifying sparse cell types in specific samples, these misclassifications suggest that there is still considerable room for improvement.

**Figure 4:**
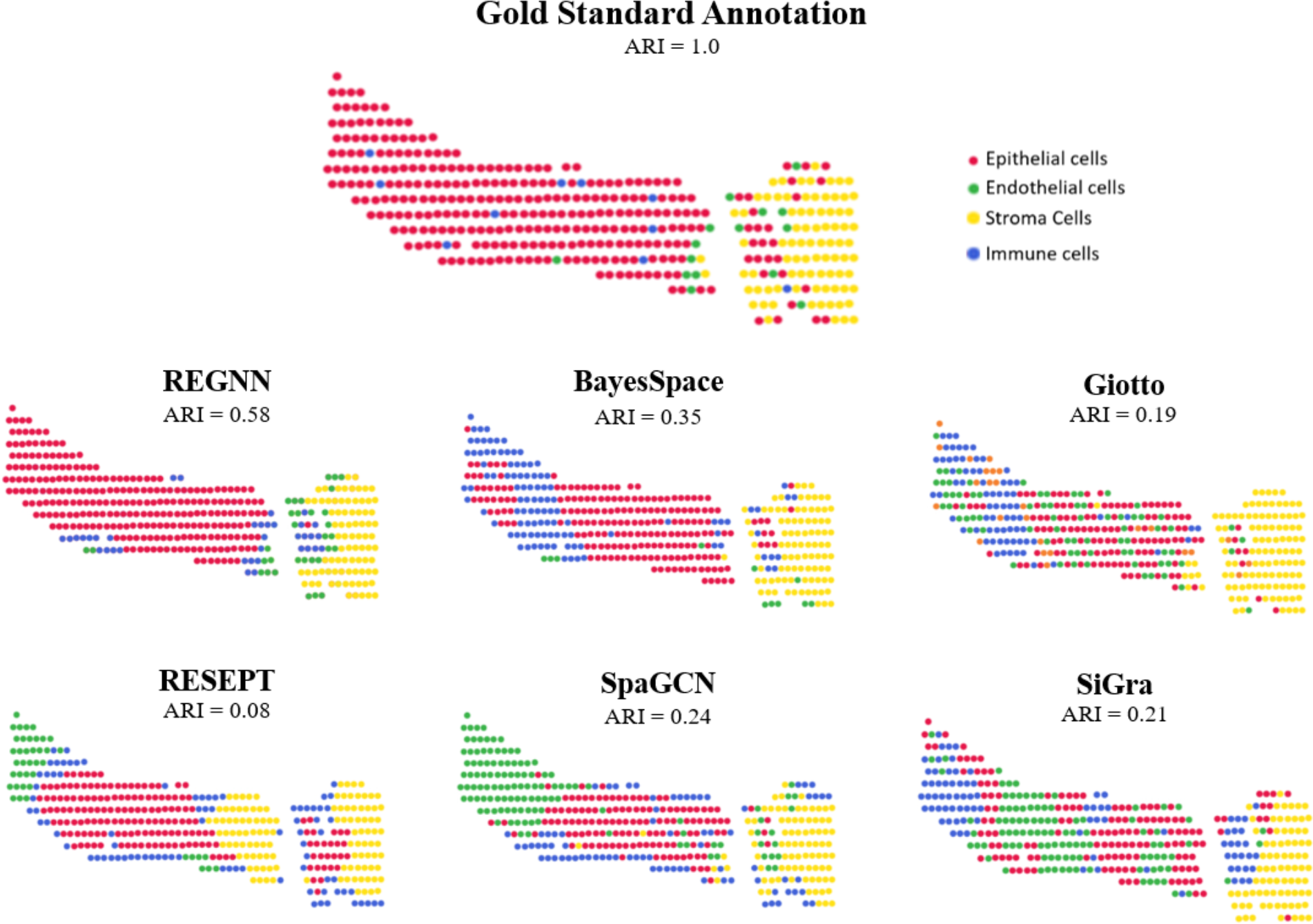
Visualization of results from computational methods. The gold standard annotations and calculated results of the computational methods are mapped to the original locations of CKD sample V10S14-085_XY04_21-0057.

### 3.2 Equivariance and PE both contribute to the expressive power of REGNN

We then investigated whether the designed equivariance and PE indeed contributed to the performances of REGNN using an ablation test. We utilized a vanilla GNN, which kept a majority of REGNN model but removed both equivariance in AGG operation and PE in UPDATE operation. Then we only kept equivariance in AGG operation and only kept PE in UPDATE operation. The results of these simplified models were compared with the REGNN model equipped with both components on the representative CKD sample V10S14-085_XY04_21-0057. From **Table 2**, we can see that directly utilizing PE improved the performance in ARI. While it did not show improvement when only incorporating equivariance, combining both equivariance and PE significantly enhanced the expressive power of REGNN. These results were consistent with the theoretical analysis of graph deep learning model, where implementing equivariance increases the expressive power through enhancing the GNN architecture, and PE increases expressive power by strengthening the topology of the GNN[28].

**Table 2:** Ablation tests with representative CKD sample V10S14-085_XY04_21-0057 - CKD.

| Ablation Tests | ARI score |
| --- | --- |
| Vanilla GNN (No EGNN or PE) | 0.3419 |
| Equivariance only (No PE) | 0.3095 |
| PE only (No equivariance) | 0.3601 |
| EGNN and PE (REGNN) | **0.5814** |

### 3.3 Performance of REGNN representation is not sensitive to clustering algorithms

Furthermore, we explored whether the implemented clustering algorithm, other than the GNN model, played a significant role in the efficacy of the results. Besides utilizing k-means in REGNN, three other clustering algorithms, including affinity propagation, agglomerative, and spectral clustering, were tested on clustering graph embedding from REGNN. From the ARI results on the representative CKD sample V10S14-085_XY04_21-0057 in **Table 3**, the performances of each clustering algorithm were very close, with K-means seeming to be the most effective clustering method. These results demonstrated the excellence of REGNN’s expressive power, where the learned embedding clearly represents relations preserved in the learned presentation, which can be easily detected and captured by various clustering algorithms.

**Table 3:**
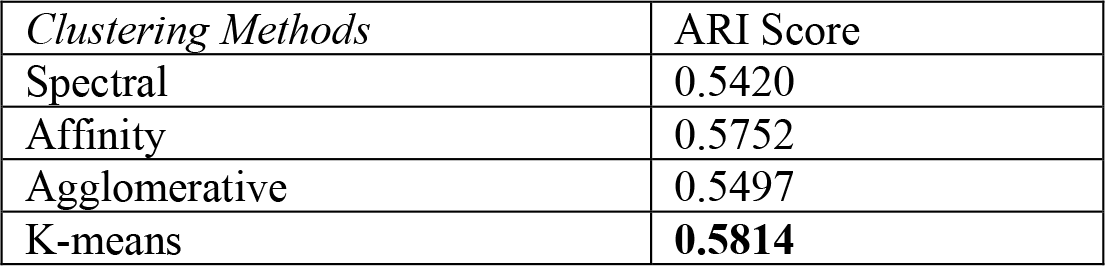
Performance of different clustering algorithms on REGNN graph embeddings on representative CKD sample V10S14-085_XY04_21-0057 - CKD.

## 4 Discussion

Graph neural networks are powerful deep learning models on graph data structures, but their inherent expressive power is theoretically limited by their capacities in modeling heterogeneous, symmetric lattices of SRT. We introduce REGNN, an expressive power enhanced graph deep learning framework designed specifically for modeling SRT data on highly heterogeneous tissues, such as samples from kidney disease. Compared with several existing computational methods developing on ribbon-like less heterogeneous brain cortex tissue, REGNN targets the more challenging mosaic-like highly heterogeneous tissue by integrating equivariance and positional encoding. In the study of SRT on kidney research, REGNN outperforms existing approaches with enhanced expressive power.

The increased expressive power of REGNN depends on the following two enhancements in GNN’s message passing mechanism: (1) Integrating the equivariance in the aggregation operation, which enables the modeling of the SRT lattice in rotational and translational symmetries. These innate symmetries bring similar topology in graph modeling, which confuses classical GNN models, and existing computational methods often omit them. (2) Integrating PE to model the relative positions of the nodes in the SRT lattice in the update operation. Like Transformer’s architecture, PE explicitly marked relative spatial relations between the nodes, which helps the model identify the nodes’ properties in the heterogenous biological context.

Although REGNN framework achieved some success in kidney studies, there are still limitations in the proposed model. (1) The current model is built on lattice from the sequencing based SRT, mostly from the 10X Visium platform. We will continue working on different image based SRT, such as FISH technologies. (2) Current improvements in resolution bring more challenges in graph modeling. Compared to the spots in 10X Visium data with thousands of nodes in the modeled graph, other advanced technologies such as MERFISH, 10X Xenium, and NanoString CosMx have hundreds of thousands of nodes in the modeled graph, which brings challenges in the scaling of the graph model. (3) Admittedly, accurately inferring the correct architecture on some kidney samples is challenging due to its intrinsic, highly heterogeneous nature. Even REGNN obtained some success in several samples, it is still an open problem in the field.

In the future, we will continue improving the REGNN’s expressive power with more advanced positional encoding[33], other high-order technologies like Mixhop[34], and incorporating other modalities such as H&E images. We are also interested in exploring cutting-edge large language models like LLaMa[30] to model the complex relations in the SRT data. Biologically, we will continue improving the model’s performance by targeting sub cell-types within the kidney, which would aid in working on downstream analyses of the differences in identified molecular profiles between different types of kidney diseases. We will also check their generalizability in other datasets from other independent data sources, such as HUPMAP[35] and HCA[36]. As we continue to fine-tune REGNN’s model on kidney tissue architecture, we aim to expand our framework’s scope so that it can robustly perform analysis on other highly heterogeneous mosaic-like tissues such as lymph nodes and colon.

## References

1. Murray, I.V. and M.A. Paolini, Histology, Kidney and Glomerulus, in StatPearls. 2023: Treasure Island (FL) ineligible companies. Disclosure: Michael Paolini declares no relevant financial relationships with ineligible companies.

2. Johansen, K.L., et al., US Renal Data System 2020 Annual Data Report: Epidemiology of Kidney Disease in the United States. Am J Kidney Dis, 2021. 77(4 Suppl 1): p. A7–A8.

3. Safari, S., et al., Epidemiology and Outcome of Patients with Acute Kidney Injury in Emergency Department; a Cross-Sectional Study. Emerg (Tehran), 2018. 6(1): p. e30.

4. Melo Ferreira, R., et al., Integration of spatial and single-cell transcriptomics localizes epithelial cell-immune cross-talk in kidney injury. JCI Insight, 2021. 6(12).

5. Moses, L. and L. Pachter, Museum of spatial transcriptomics. Nat Methods, 2022. 19(5): p. 534–546.

6. Method of the Year 2020: spatially resolved transcriptomics. Nat Methods, 2021. 18(1): p. 1.

7. Rao, A., et al., Exploring tissue architecture using spatial transcriptomics. Nature, 2021. 596(7871): p. 211–220.

8. Jin, S., et al., Inference and analysis of cell-cell communication using CellChat. Nat Commun, 2021. 12(1): p. 1088.

9. Svensson, V., S.A. Teichmann, and O. Stegle, SpatialDE: identification of spatially variable genes. Nat Methods, 2018. 15(5): p. 343–346.

10. Wang, J., et al., Dimension-agnostic and granularity-based spatially variable gene identification. Res Sq, 2023.

11. Elosua-Bayes, M., et al., SPOTlight: seeded NMF regression to deconvolute spatial transcriptomics spots with single-cell transcriptomes. Nucleic Acids Res, 2021. 49(9): p. e50.

12. Zhao, E., et al., Spatial transcriptomics at subspot resolution with BayesSpace. Nat Biotechnol, 2021. 39(11): p. 1375–1384.

13. Dries, R., et al., Giotto: a toolbox for integrative analysis and visualization of spatial expression data. Genome Biol, 2021. 22(1): p. 78.

14. Wang, J., et al., scGNN is a novel graph neural network framework for single-cell RNA-Seq analyses. Nat Commun, 2021. 12(1): p. 1882.

15. Hu, J., et al., SpaGCN: Integrating gene expression, spatial location and histology to identify spatial domains and spatially variable genes by graph convolutional network. Nat Methods, 2021. 18(11): p. 1342–1351.

16. Chang, Y., et al., Define and visualize pathological architectures of human tissues from spatially resolved transcriptomics using deep learning. Comput Struct Biotechnol J, 2022. 20: p. 4600–4617.

17. He, K., et al. Deep Residual Learning for Image Recognition. in 2016 IEEE Conference on Computer Vision and Pattern Recognition (CVPR). 2016.

18. Tang, Z., et al., SiGra: single-cell spatial elucidation through an image-augmented graph transformer. Nat Commun, 2023. 14(1): p. 5618.

19. Lake, B.B., et al., An atlas of healthy and injured cell states and niches in the human kidney. Nature, 2023. 619(7970): p. 585–594.

20. Kipf, T.N. and M. Welling, Semi-Supervised Classification with Graph Convolutional Networks. arXiv [cs.LG], 2017.

21. Xu, K., et al., How Powerful are Graph Neural Networks? arXiv [cs.LG], 2019.

22. Balcilar, M., et al., Breaking the Limits of Message Passing Graph Neural Networks, in Proceedings of the 38th International Conference on Machine Learning, M. Marina and Z. Tong, Editors. 2021, PMLR: Proceedings of Machine Learning Research. p. 599--608.

23. Elmarakeby, H.A., et al., Biologically informed deep neural network for prostate cancer discovery. Nature, 2021. 598(7880): p. 348–352.

24. de Boer, I.H., et al., Rationale and design of the Kidney Precision Medicine Project. Kidney Int, 2021. 99(3): p. 498–510.

25. The cerebral cortex, meninges, basal ganglia, and ventricular system - Knowledge @ AMBOSS.

26. Maynard, K.R., et al., Transcriptome-scale spatial gene expression in the human dorsolateral prefrontal cortex. Nat Neurosci, 2021. 24(3): p. 425–436.

27. Satorras, V.G., E. Hoogeboom, and M. Welling, E(n) Equivariant Graph Neural Networks. arXiv [cs.LG], 2022.

28. Sato, R., A Survey on The Expressive Power of Graph Neural Networks. arXiv [cs.LG], 2020.

29. Vaswani, A., et al., Attention is All you Need, in Advances in Neural Information Processing Systems, I. Guyon, et al., Editors. 2017, Curran Associates, Inc. p. 5998–6008.

30. Touvron, H., et al., LLaMA: Open and Efficient Foundation Language Models. arXiv [cs.CL], 2023.

31. Zhong, E.D., et al., CryoDRGN: reconstruction of heterogeneous cryo-EM structures using neural networks. Nat Methods, 2021. 18(2): p. 176–185.

32. Rodriguez, M.Z., et al., Clustering algorithms: A comparative approach. PLoS One, 2019. 14(1): p. e0210236.

33. Ke, G., D. He, and T.-Y. Liu, Rethinking Positional Encoding in Language Pre-training. arXiv [cs.CL], 2021.

34. Abu-El-Haija, S., et al., MixHop: Higher-Order Graph Convolutional Architectures via Sparsified Neighborhood Mixing. arXiv [cs.LG], 2019.

35. Snyder, M.P., et al., The human body at cellular resolution: the NIH Human Biomolecular Atlas Program. Nature, 2019. 574(7777): p. 187–192.

36. Regev, A., et al., The Human Cell Atlas. Elife, 2017. 6.

